# Comparing the electrophysiology and morphology of human and mouse layer 2/3 pyramidal neurons with Bayesian networks

**DOI:** 10.1101/2020.06.02.130252

**Authors:** Bojan Mihaljević, Pedro Larrañaga, Concha Bielza

## Abstract

Pyramidal neurons are the most common neurons in the cerebral cortex. Understanding how they differ between species is a key challenge in neuroscience. We compared human temporal cortex and mouse visual cortex pyramidal neurons from the Allen Cell Types Database in terms of their electrophysiology and basal dendrites’ morphology. We found that, among other differences, human pyramidal neurons had a higher threshold voltage, a lower input resistance, and a larger basal dendritic arbor. We learned Gaussian Bayesian networks from the data in order to identify correlations and conditional independencies between the variables and compare them between the species. We found strong correlations between electrophysiological and morphological variables in both species. One result is that, in human cells, dendritic arbor width had the strongest effect on input resistance after accounting for the remaining variables. Electrophysiological variables were correlated, in both species, even with morphological variables that are not directly related to dendritic arbor size or diameter, such as mean bifurcation angle and mean branch tortuosity. Contrary to previous results, cortical depth was correlated with both electrophysiological and morphological variables, and its effect on electrophysiological could not be explained in terms of the morphological variables. Overall, the correlations among the variables differed strikingly between human and mouse neurons. Besides identifying correlations and conditional independencies, the learned Bayesian networks might be useful for probabilistic reasoning regarding the morphology and electrophysiology of pyramidal neurons.

## 1 Introduction

A key challenge in neuroscience is to understand how pyramidal neurons differ across species and cortical regions^1–7^. They are often compared in terms of their dendritic morphology, since it directly influences neuronal computation^8–10^. Compared to rodents, human pyramidal neurons’ dendrites are larger^4^ and have more synaptic connections per cell^11, 12^, while the human cortical layer 2/3 is thicker^3, 12^, with easily distinguished layers 2 and 3. Dendritic morphology also varies across cortical regions within a single species^4, 7, 13–16^, although more so in primates than in rodents^1, 2, 17^. For example, the pyramidal neurons of the monkey visual cortex have smaller basal dendrites than those of its prefrontal cortex, while there is no significant difference in the mouse^2^. In terms of electrophysiology, mouse visual cortex pyramidal neurons have a lower threshold voltage, shorter rise time and longer fall time than than those of rhesus monkey, while there is no significant difference in subthreshold features such as time constant and input resistance^2^.

There has been comparatively little analysis of how the different electrophysiological and morphological variables correlate with each other and how do these correlations vary between species. In particular, there has been limited study of correlation between electrophysiology and morphological variables. Exceptions are Ref.^2^, which found that larger neurons had a lower input resistance, and Ref.^18^, which found that action potential is accelerated in neurons with larger dendritic surface area. Also Ref.^16^ found that the morphology of human pyramidal neurons varied with cortical depth while the electrophysiology did not. These analyses were limited to estimating the linear correlation between pairs of variables. This ignores the effects of confounders as well as the conditional independencies among variables. Instead, dependencies can be quantified with partial correlation coefficients and modelled with graphical models^19^. One type of graphical models that is useful for modeling conditional independencies are Bayesian networks^20, 21^. These models, based on directed acyclic graphs, let us visualise the probabilistic relationships between the variables. Their applications in neuroscience^22, 23^ include interneuron classification^24, 25^ and the generation of synthetic dendritic branches^26^.

In this paper, we compare layer 2/3 human temporal cortex and mouse visual cortex pyramidal neurons from the Allen Cell Types Database in terms of their electrophysiology and basal dendrites’ morphology. The Allen Cell Type Database cells are unique in that they have both electrophysiology and morphology data and these data were obtained with a standardized procedure for both species. Since some cell’s apical dendrites were incompletely reconstructed, we only study the morphological features of the basal dendrites. We learn from data Gaussian Bayesian networks in order to identify correlations and conditional independencies between the variables. We learn Bayesian networks from three different subsets of our data set: (a) from electrophysiological variables alone; (b) from morphological variables alone; and (c) from electrophysiological and morphological variables combined. For each data subset we learn one Bayesian network per species, which yields a total of six networks.

The rest of this paper is structured as follows. Section 2 describes the data set, the variables and analysis methodology. Section 3 provides the results. We discuss our findings in Section 4.

## 2 Methods

### 2.1 Data

We used adult human and adult mouse neurons from the Allen Cell Type Database (http://celltypes.brain-map.org/). Human cells were acquired from donated ex vivo brain tissue. We used all excitatory (spiny) cells from layers 2 and 3 of the temporal (human) and visual (mouse) cortex that had a reconstructed morphology. Our sample consisted of 42 human cells from the temporal cortex and 21 mouse cells from the visual cortex.

### 2.2 Electrophysiological variables

The Allen Cell Type Database provides pre-computed electrophysiological features. These features were derived from high temporal resolution data on membrane potential measurements (in current-clamp mode) obtained with a standardized patch clamp protocol. We used 11 electrophysiological features provided by Allen Cell Type Database, covering subthreshold and suprathreshold features of the cells, including those relative to action potentials. Below we list these features along with brief descriptions (see also Table 1 for their mean values) while we refer the reader to the technical white paper by the Allen Cell Type Database for details (http://help.brain-map.org/download/attachments/8323525/CellTypes_Ephys_Overview.pdf).

**Table 1.** Per-species mean ± standard deviation for each electrophysiological variable, along with the p-value of the t-test.

| Variable | Human | Mouse | p-value |
| --- | --- | --- | --- |
| rest (mV) | -71.77 ± 3.75 | -77.69 ± 4.06 | 0.00 |
| resistance (M Ω) | 83.96 ± 54.31 | 151.61 ± 68.86 | 0.00 |
| tau (ms) | 27.43 ± 9.01 | 20.02 ± 9.87 | 0.01 |
| threshold (pA) | 203.33 ± 100.21 | 186.67 ± 115.94 | 0.58 |
| peak (mV) | 46.31 ± 4.77 | 37.30 ± 8.68 | 0.00 |
| amplitude (mV) | 99.36 ± 7.35 | 91.07 ± 9.07 | 0.00 |
| up down ratio | 3.65 ± 0.81 | 3.70 ± 0.83 | 0.82 |
| rise time (ms) | 0.00 ± 0.00 | 0.00 ± 0.00 | 0.00 |
| fall time (ms) | 0.08 ± 0.17 | 0.14 ± 0.28 | 0.35 |
| latency (ms) | 0.08 ± 0.03 | 0.06 ± 0.02 | 0.00 |
| f-i curve () | 0.08 ± 0.05 | 0.14 ± 0.08 | 0.00 |

Subthreshold features were computed as follows. Resting potential (`rest`): Average pre-stimulus membrane potential across all the long square responses; Input resistance (`resistance`): The slope of a linear fit of minimum membrane potentials during the responses onto their respective stimulus amplitudes, for long square sweeps with negative current amplitudes that did not exceed 100 pA; Time constant (`tau`): Exponential curve fit between 10% of the maximum voltage deflection (in the hyperpolarizing direction) and the minimum membrane potential during the response, the time constants of these fits were averaged across steps to estimate the membrane of the cell.

All action potential waveforms were evoked by a long square (one second) current step stimulus. The waveforms of the first action potentials were collected from each cell and aligned on the time of their thresholds. Action potential features were computed as follows. Threshold (`threshold`): The level of injected current at threshold; Peak (`peak`): Maximum value of the membrane potential during the action potential; Amplitude (`amplitude`): Difference between the action potential trough and the action potential peak, where the trough is the minimum value of the membrane potential between the peak and the next action potential; Upstroke/downstroke ratio (`up down ratio`): The ratio between the absolute values of the action potential peak upstroke and the action potential peak downstroke, where the upstroke is the maximum value of dV/dt between the threshold and the peak, and peak downstroke is the minimum value of dV/dt between the peak and the trough; Rise time (`rise time`): Time from threshold to the peak; fall time (`fall time`): Time from peak to the trough.

Additional suprathreshold features were computed as follows. Latency (`latency`): Time between the start of the stimulus until the first spike; “f-i curve” (`f-i curve`): Slope of a straight line fit to the suprathreshold part of the curve of frequency response of the cell versus stimulus intensity for long square responses.

### 2.3 Morphological variables

The Allen Cell Type Database provides 3D neuron morphology reconstructions. These were obtained by filling the cells with biocytin and serially imaged to visualize their morphologies. Detailed description of the reconstruction protocol is provided in the Allen Cell Type Database morphology overview technical whitepaper (http://help.brain-map.org/download/attachments/8323525/CellTypes_Morph_Overview.pdf).

Because the apical trees of some neurons were incompletely reconstructed, we only computed morphological features of the basal dendrites. We computed 8 features using the open-source NeuroSTR library (https://computationalintelligencegroup.github.io/neurostr/). Below we list these features along with brief descriptions (see also Table 2 for their mean values).

**Table 2.** Per-species mean ± standard deviation for each morphological variable, along with the p-value of the t-test.

| Variable | Human | Mouse | p-value |
| --- | --- | --- | --- |
| `distance` ($\mu m$) | 133.00 ± 63.00 | 63.00 ± 5.00 | 0.00 |
| `tortuosity` | 1.15 ± 0.05 | 1.14 ± 0.05 | 0.52 |
| `angle` (rad) | 1.01 ± 0.14 | 1.13 ± 0.18 | 0.01 |
| `diameter` ($\mu m$) | 0.33 ± 0.12 | 0.27 ± 0.06 | 0.01 |
| `height` ($\mu m$) | 403.00 ± 262.00 | 192.00 ± 22.00 | 0.00 |
| `width` ($\mu m$) | 423.00 ± 147.00 | 227.00 ± 48.00 | 0.00 |
| `depth` ($\mu m$) | 114.00 ± 26.00 | 62.00 ± 17.00 | 0.00 |
| `total_length` ($\mu m$) | 5008.00 ± 3108.00 | 1867.00 ± 540.00 | 0.00 |

A number of features were averaged across all bifurcations or branches of the basal dendrites and were computed as follows. Average path distance (`distance`): Sum of the lengths of all compartments’ length starting from the dendrites’ insertion point into the soma up the bifurcation point, averaged over all bifurcation points; Average branch tortuosity (`tortuosity`): Ratio of branch length and the length of the straight line between the beginning and the end of a branch, averaged over all branches; Average remote bifurcation angle (`angle`): Shortest planar angle between the vectors from the bifurcation to the endings of the daughter branches, averaged across all bifurcations; Average branch diameter (`diameter`).

Global basal dendrites features were computed as follows. Height (`height`): Difference between the maximum and minimum values of Y-coordinates of the dendrites; Width (`width`): Difference between the maximum and minimum values of X-coordinates of the dendrites; Depth (`depth`): Difference between the maximum and minimum values of Z-coordinates of the dendrites; Total length (`totallength`): Sum of branch length of all the branches of the dendrites.

The Allen Cell Type Database also provided the depth of each cell’s soma (`rel depth`) relative to pia and white matter.

### 2.4 Bayesian networks

A Bayesian network (BN)^21^ *ℬ* allows us to compactly encode a joint probability distribution over a vector of *n* random variables **X** by exploiting conditional independencies among triplets of sets of variables in **X** (e.g., *X* is independent of *Y* given *Z*). A BN consists of a directed acyclic graph (DAG) *𝒢* and a set of parameters ***θ*** (*ℬ* = (*𝒢*, ***θ***)). The vertices (i.e., nodes) of *𝒢* correspond to the variables in **X** while its directed edges (i.e., arcs) encode the conditional independencies among **X**. A joint probability density *f*_*𝒢*_ (**x**) encoded by *ℬ*, where **x** is an assignment to **X**, factorizes as a product of local conditional densities,

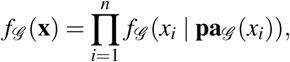

where **pa**_*𝒢*_ (*x*_*i*_) is an assignment to variables **Pa**_*𝒢*_ (*X*_*i*_), the set of parents of *X*_*i*_ in **X** according to *𝒢*. *𝒢* induces conditional independence constraints for *f*_*𝒢*_ (·), derivable from the basic constraints that each *X*_*i*_ is independent of its non-descendents in *𝒢* given **Pa**_*𝒢*_ (*X*_*i*_). For example, for any pair of variables *X,Y* in **X** that are not connected by an arc in *𝒢* there exists a set of variables **Z** in **X** (disjoint from *{X}* and *{Y}*) such that *X* and *Y* are independent conditionally on **Z** (i.e., *f*_*𝒢*_ (*X,Y* |**Z**) = *f*_*𝒢*_ (*X* | **Z**) *f*_*𝒢*_ (*Y* | **Z**)). Similarly, for any pair of variables *X,Y* in **X** that are connected by an arc in *𝒢* there is no set **Z** such that *X* and *Y* are independent conditionally on **Z**. These constraints extend to nodes not connected by an arc in *𝒢* and the structure *𝒢* thus lets us identify conditional independence relationships among any triplet of sets of variables *X, Y*, and **Z** in **X**. For example, in the DAG *X → Y → Z* we only have one independence: *X* is independent of *Z* conditional on *Y*; *X* and *Y, X* and *Z*, and *Y* and *Z* are not marginally independent. The Markov blanket of *X*_*i*_ is the set of variables **MB**(*X*_*i*_) such that *X*_*i*_ is independent of **X** *\* **MB**(*X*_*i*_) conditional on **MB**(*X*_*i*_). The Markov blanket of *X*_*i*_ is easily determined from *𝒢* as it corresponds to the parents, the children and the spouses (other parents of the children of *X*_*i*_) of *X*_*i*_ in *𝒢*.

The parameters ***θ*** specify the local conditional densities *f*_*𝒢*_ (*x*_*i*_ | **pa**_*𝒢*_ (*x*_*i*_)) for each variable *X*_*i*_. When **X** contains only continuous variables, as in our case, a common approach is to let *f*_*𝒢*_ (**x**) be a multivariate normal density. The local conditional density for *X*_*i*_ is 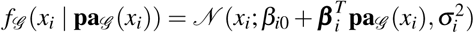. There is thus a different vector of coefficients 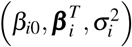 for each *X*_*i*_.

### 2.5 Learning Bayesian networks from data

Learning a Bayesian network *ℬ* from a data set *𝒟* = {**x**^1^,*…*, **x**^*N*^} of *N* observations of **X** involves two steps: (a) learning the DAG *𝒢*; and (b) learning ***θ***, the parameters of the local conditional distributions. There are two main approaches to learning *𝒢* from *𝒟* ^21^: (a) by testing for conditional independence among triplets of sets of variables (the *constraint-based* approach); and (b) by searching the space of DAGs in order to optimize a score such as penalized likelihood (the *score-based* approach). While seemingly very different, conditional independence tests and network scores are related statistical criteria^27^. For example, when considering whether to include the arc *Y → X* into a graph *𝒢*, the likelihood-ratio test of conditional independence of *X* and *Y* given **Pa**_*𝒢*_ (*X*) and the Bayesian information criterion^28^ (BIC) score are both functions of log 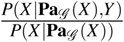. They differ in computing the threshold for determining independence: the former relies on the distribution of the statistic under the null model (i.e., conditional independence) whereas the latter is based on an approximation to the Bayes factor between the null and alternative models. Besides using different criteria, the constraint-based and score-based approaches also differ in model search, that is, in terms of the sets *X, Y*, and **Z** that they choose to test conditional independence for. The score-based approaches tend to be more robust^21^, as they may reconsider previous steps in the search by removing or reversing previously added arcs. We thus followed a score-based approach in this paper.

### 2.6 Settings

We learned network structures by using the tabu algorithm^29^, implemented in the bnlearn^30^ R^31^ package, to optimize the BIC score. The tabu algorithm is a local search that efficiently allows for score-degrading operators by avoiding those that undo the effect of recently applied operators; we used a tabu list of size 30 and allowed for up to 30 iterations without improving network score.

## 3 Results

We first look at electrophysiological (Section 3.1) and morphological features (Section 3.2) separately, and then learn joint Bayesian networks for both electrophysiological and morphological features (Section 3.3).

### 3.1 Electrophysiology

All variables except for `threshold`, `up down ratio`, and `fall time` differed significantly between the species (Table 1). Human neurons had lower a `resistance`, higher time constant (`tau`), `rest`, `peak` voltage, `amplitude` and `latency`, a longer action potential `rise time` and a shorter `fall time`.

The human and mouse BNs uncovered relevant correlations and independencies among the variables. In the human BN (Figure 1a), `rel depth` was correlated with `rest`, `threshold`, `up down ratio`, `latency`, `f-i curve` and `rel depth` while its Markov blanket consisted of `tau`, `threshold`, `peak`, `up down ratio` and `f-i curve`. This is contrary to Ref.^16^ which found that human electrophysiological features such as input resistance and membrane time constant were independent of depth in the human L2/3 pyramidal neurons of the temporal cortex. In particular, `rel depth` had a strong positive partial correlation with `up down ratio` and a strong negative one with `threshold`. The `fall time` was uncorrelated with other variables, `rise time` was independent of other variables given `amplitude`, while `rest` was independent of other variables given `up down ratio` and `threshold`. All other variables had Markov blankets of size three or larger, with the largest being that of `up down ratio`, with seven variables. The strongest partial correlation was that between `peak` and `amplitude`. See Figure 1a for all partial correlation coefficients.

**Figure 1.**
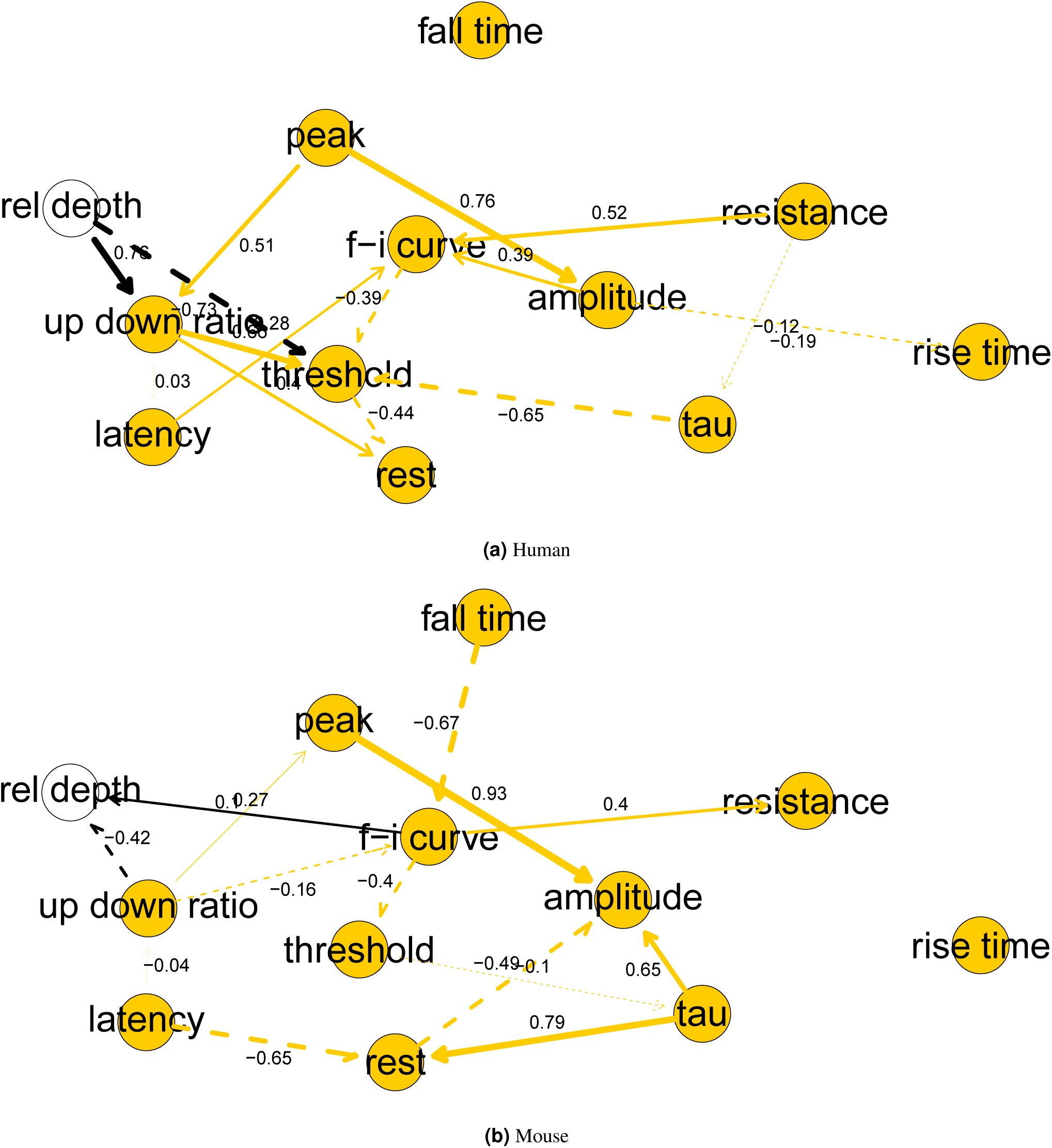
Bayesian networks for electrophysiological features. Arc width is proportional to the absolute value of the partial correlation (shown above the arc) between the nodes. Arcs corresponding to negative partial correlations plotted with dashed lines. Proximity between two nodes is unrelated to the magnitude of partial correlation.

In the mouse BN (Figure 1b), `rel depth` was only correlated with `up down ratio` and `f-i curve`, the two variables which formed its Markov blanket while, contrary to the human BN, its partial correlation with `up down ratio` was negative. `rise time` was uncorrelated with other variables, `fall time` was independent of other variables given `f-i curve`, while `rest` was independent of other variables given `latency` and `tau`. The remaining variables had Markov blankets of size three or larger, with the largest being those of `up down ratio` and `f-i curve`, with seven variables. The strongest partial correlations were those between `peak` and `amplitude`, and `tau` and `rest`. See Figure 1b for all partial correlation coefficients.

Overall, the human and mouse BNs were strikingly different, with only two common arcs (`peak` *→* `amplitude` and `f-i curve` *→* `threshold`). No variable had an identical Markov blanket in the two graphs. While the magnitudes of `threshold`, `fall time` and `up down ratio` did not differ significantly between the species (Table 1), the BNs show that their correlation with other variables did. A rare common feature of the two BNs was the strong positive partial correlation between `amplitude` and `peak`.

### 3.2 Morphology

All variables except for `tortuosity` differed significantly between the species (Table 2). Human basal neurons were larger, had thicker branches and sharper bifurcation angles. The 2.7-fold difference in total length was more pronounced than the 2.1-fold difference that Ref.^4^ observed for basal dendrites of human and mouse temporal cortex pyramidal neurons.

In the human BN (Figure 2a), `rel depth` was marginally correlated with all variables except for `angle` and `depth`. However, it was independent of all other variables given `tortuosity`, with deeper neurons being more straight (i.e., having lower `tortuosity` values), yet with virtually no effect on overall dendritic size (correlation coefficient, *ρ* = 0.03). This is also contrary to the results of Ref.^16^, which found a correlation coefficient *ρ* = 0.50 between basal dendrites total length and depth from the pia. An interesting partial correlation was that between `diameter` and `tortuosity`, indicating that cells with thicker branches tended to have more tortuous branches; this is surprising because thicker branches tend to be shorter and one might expect shorter branches to be less tortuous.

**Figure 2.**
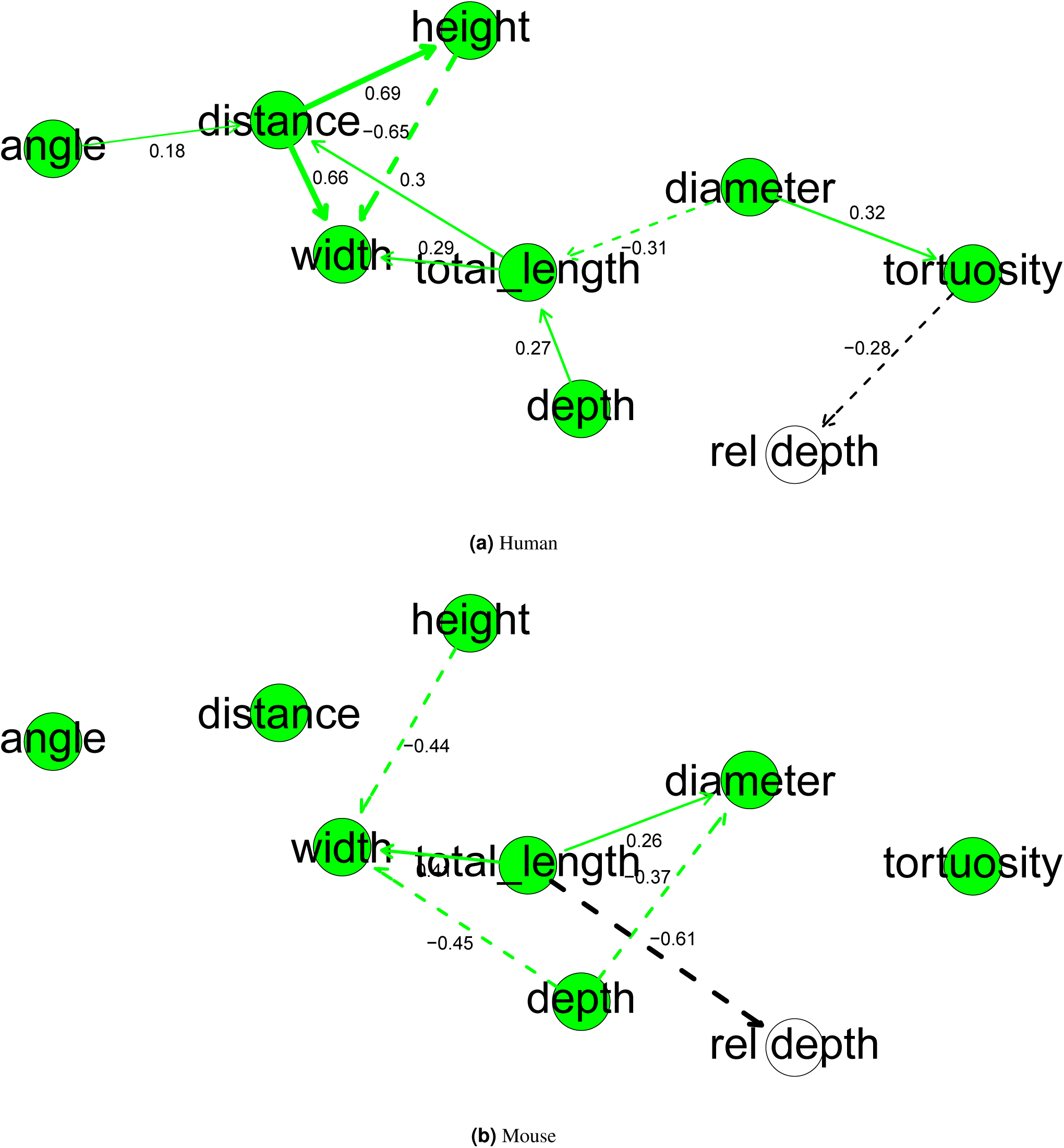
Bayesian networks for morphological features. Arc width is proportional to the absolute value of the partial correlation (shown above the arc) between the nodes. Arcs corresponding to negative partial correlations plotted with dashed lines. Proximity between two nodes is unrelated to the magnitude of partial correlation.

In the mouse BN (Figure 2b), `rel depth` had a strong negative partial correlation with `totallength`. Their marginal (not partial) correlation was also negative (−0.73), meaning that deeper cells had smaller basal dendrites. This is the opposite of the effect that Ref.^16^ observed for human neurons and contrary to what observed on temporal cortex mouse cells, where they found no significant change in morphological features with increasing depth. Unlike in the human, cells with larger basal dendrites tended to have thicker dendrites (*ρ* = 0.43) while `tortuosity` was independent of all other variables.

Overall, the human and mouse BNs were strikingly different, with only two common arcs (`totallength` *→* `width` and `height` *→* `width`). No variable had an identical Markov blanket in the two graphs.

### 3.3 Electrophysiology and morphology

The human BN (Figure 3a) shows that electrophysiological and morphological variables were highly correlated. The BN had 10 arcs between electrophysiological and morphological features and, except for `latency` and `fall time`, all electrophysi-ological features had at least one morphological variable in their Markov blankets, with `depth` being the only morphological variable not present in these Markov blankets. In particular, `resistance` had arcs to four morphological features and seven morphological variables in its Markov blanket. While it was already known that `resistance` decreases with dendritic size^2^, we found that, after accounting for `totallength` and the remaining variables, the strongest effect was that of dendritic arbor `width`. We also found that `peak` increased with `totallength` and decreased with `tortuosity`, both after accounting for `totallength` and marginally (*ρ* = *−*0.37), while `angle` had positive partial correlations with `resistance` and `rise` `time`. Conversely to the above-stated, `depth` was the only morphological variable without electrophysiological features in its Markov blanket while `fall time` and `latency` were absent from the Markov blankets of morphological features. The Markov blanket of `height` included eight electrophysiological features. As in Figure 1a and Figure 2a, `rel depth` was directly correlated with `up down ratio`, `threshold` and `tortuosity`. Figure 3a also shows that the correlation of `rel depth` with the electrophysiological variables cannot be explained as an indirect effect via differences in morphology with respect to cortical depth, as `rel depth` is not independent of electrophysiological variables given the morphological ones. Instead, it corresponds to intrinsic differences in electrophysiology with regards to cortical depth that are unexplained with the used morphological variables. This was already suggested by Figure 1a and Figure 2a, which showed a stronger correlation of `rel depth` with electrophysiological than with morphological variables.

**Figure 3.**
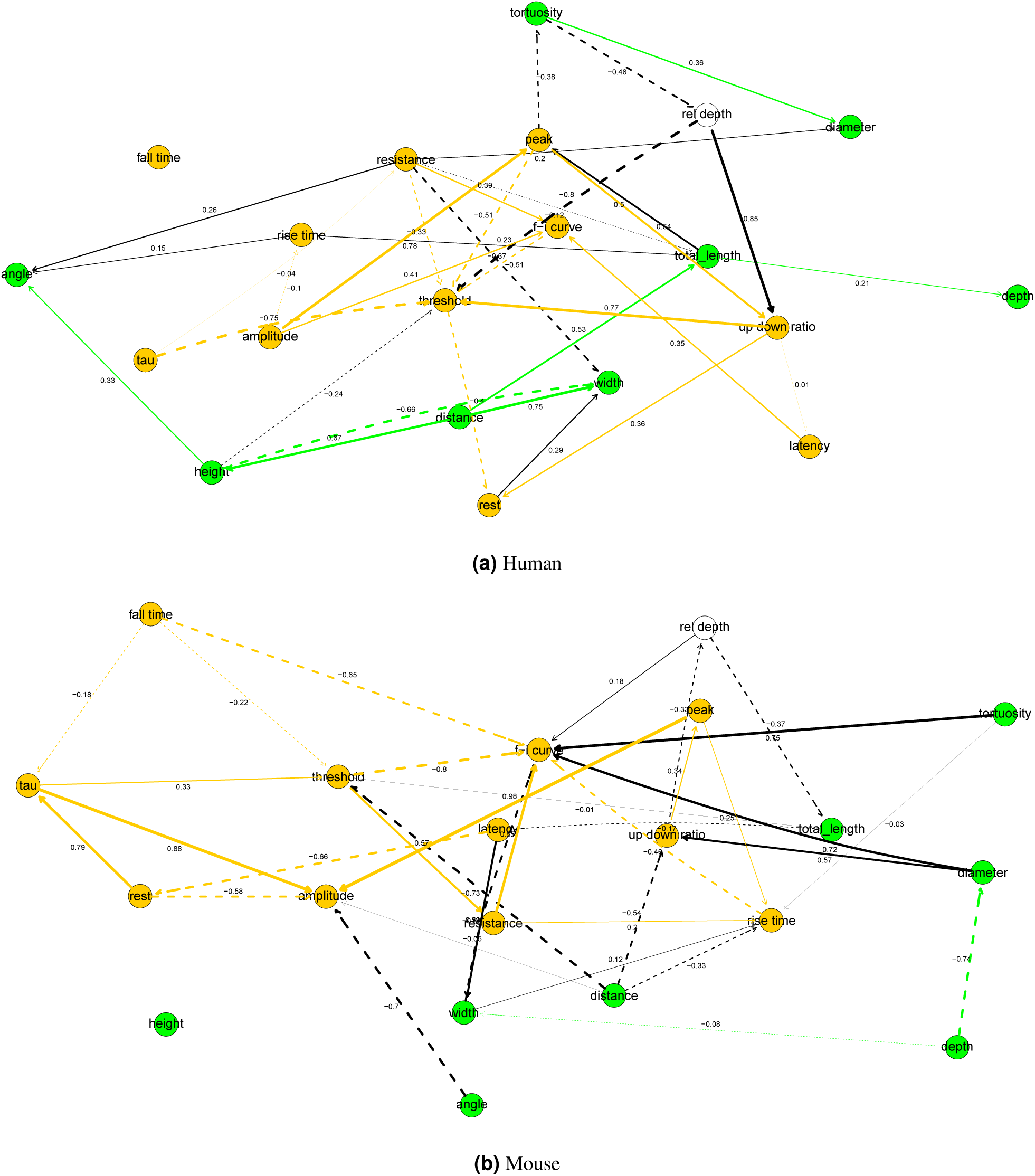
Bayesian networks for electrophysiological and morphological features. Electrophysiological nodes and the arcs between them shown in green; morphological nodes and the arcs between them in orange. Arc width is proportional to the absolute value of the partial correlation (shown above the arc) between the nodes. Arcs corresponding to negative partial correlations plotted with dashed lines. Proximity between two nodes is unrelated to the magnitude of partial correlation.

The mouse BN (Figure 3b) also showed strong correlation between electrophysiological and morphological variables. The BN had 14 arcs between electrophysiological and morphological features and all electrophysiological variables had at least two morphological variables in their Markov blanket. The strongest partial correlations were those between `diameter` and `f-i curve`, `diameter` and `up down ratio`, and `tortuosity` and `f-i curve`, `angle` and `amplitude`, and `distance` and `threshold`. While the Markov blanket of `resistance` included four morphological variables (`width`, `distance`, `tortuosity` and `diameter`), it only had strong or moderate marginal (that is, not partial) correlation with `diameter` and `distance` (*ρ* = *−*0.45, and *ρ* = 0.26, respectively), with the negatively correlation with `distance` indicating that it decreases with increasing dendritic size. As in the human BN, the Markov blanket of `rel depth` included both electrophysiological and morphological variables.

Overall, the two BNs were different, with only 2 common arcs: `resistance` *→* `f-i curve` and `tau` *→* `threshold`. No variable had an identical Markov blanket in the two graphs.

## 4 Discussion

We found strong differences between the electrophysiology and morphology of human and mouse pyramidal neurons, both in terms of the variables’ magnitudes and in terms of correlations between the variables, as evidenced by the differences in their Bayesian networks.

For both species, we identified the conditional independencies among electrophysiological, morphological, as well as between electrophysiological and morphological variables. In particular, we found strong correlations between electrophysiological and morphological variables in both species. We confirmed the previous finding that input resistance decreases with dendritic size. We also found that, in human cells, after accounting for dendritic arbor length and the remaining variables, dendritic arbor width had the strongest effect on the value of input resistance. We also found novel correlations between electrophysiological and morphological in both species. In human cells, peak voltage increased with dendritic arbor length and decreased with tortuosity, while input resistance and rise time were possitively correlated with the bifurcation angle after accounting for the remaining variables. In mouse cells, strong partial correlations included those between `tortuosity` and `f-i curve`, and `angle` and `amplitude`. Thus, electrophysiological variables were also correlated with morphological variables not directly related to dendritic arbor size or diameter, such as mean bifurcation angle and mean tortuosity.

Our results regarding the effect of cortical depth on the electrophysiology and the morphology of pyramidal neurons are contrary to those from Ref.^16^. They found that, in human pyramidal neurons, the size of the dendritic basal arbor increased with cortical depth while electrophysiological features were unaffected; in mouse pyramidal neurons they found no effect in morphological features while they did not assess the effect on electrophysiological ones. We, however, found that cortical depth was correlated with electrophysiological variables in both species and with morphological ones in human cells. Interestingly, we found that dendritic basal arbor size in fact decreased in mouse cells. The effect of cortical depth differed between two species, perhaps reflecting differences in laminar organization of L2 and L3 between the two species. We also showed that the correlation with electrophysiological features could not be explained in terms of the morphological variables alone. A possible explanation for the differences in the mouse cells is that we studied visual cortex cells, while Ref.^16^ studied temporal cortex cells.

Provided that our assumption of a multivariate Gaussian distribution of the variables holds, the learned Bayesian networks can be useful beyond identifying the independencies and correlations between variables. For example, they would allow for probabilistic reasoning regarding the morphology and electrophysiology of pyramidal neurons. For example, we could set the morphological variables to particular values and study the conditional distribution of electrophysiological variables. One might also use them for multi-output regression^32^, for example to predict the values of electrophysiological variables from those of the morphological ones.

## 5 Data availability

All data are available from the Allen Cell Type Database. Analysis code is available at https://github.com/ComputationalIntelligenceGroup/species-compare-electro-morpho.

## 6 Acknowledgements

This work has been partially supported by the Spanish Ministry of Economy and Competitiveness through the TIN2016-79684-P project. This project has received funding from the European Union’s Horizon 2020 Framework Programme for Research and Innovation under the Specific Grant Agreement No. 785907 (Human Brain Project SGA2).

## 7 Author contributions statement

B.M. designed and conducted the analysis and wrote the manuscript. All authors substantially reviewed the manuscript.

## 8 Additional information

The authors declare no competing interests.

